# Metabolic model of nitrogen-fixing obligate aerobe *Azotobacter vinelandii* demonstrates adaptation to oxygen concentration and metal availability

**DOI:** 10.1101/2021.06.16.448589

**Authors:** Alexander B. Alleman, Florence Mus, John W. Peters

## Abstract

There is considerable interest in promoting biological nitrogen fixation as a mechanism to reduce the inputs of nitrogenous fertilizers in agriculture, a problem of agronomic, economic, and environmental importance. For the potential impact of biological nitrogen fixation in agriculture to be realized, there are considerable fundamental knowledge gaps that need to be addressed. Biological nitrogen fixation or the reduction of N_2_ to NH_3_ is catalyzed by nitrogenase which requires a large amount of energy in the form of ATP and low potential electrons. Nitrogen-fixing organisms that respire aerobically have an advantage in meeting the energy demands of biological nitrogen fixation but face challenges of protecting nitrogenase from inactivation in the presence of oxygen. Here, we have constructed a genome-scale metabolic model of the aerobic metabolism of nitrogen-fixing bacteria *Azotobacter vinelandii*, which uses a complex electron transport system, termed respiratory protection, to consume oxygen at a high rate keeping intracellular conditions microaerobic. Our model accurately determines growth rate under high oxygen and high substrate concentration conditions, demonstrating the large flux of energy directed to respiratory protection. While respiratory protection mechanisms compensate the energy balance in high oxygen conditions, it does not account for all substrate intake, leading to increased maintenance rates. We have also shown how *A. vinelandii* can adapt under different oxygen concentrations and metal availability by rearranging flux through the electron transport system. Accurately determining the energy balance in a genome-scale metabolic model is required for future engineering approaches.

**Importance:** The world’s dependence on industrially produced nitrogenous fertilizers has created a dichotomy of issues. Some parts of the globe lack access to fertilizers and associated poor crop yields, significantly limiting nutrition, contributing to disease and starvation. In contrast, in other parts of the world, abundant nitrogenous fertilizers and associated overuse result in compromised soil quality and downstream environmental issues. There is considerable interest in expanding the impacts of biological nitrogen fixation to promote improved crop yields in places struggling with access to industrial fertilizers and reducing fertilizers’ inputs in areas where overuse is resulting in the degradation of soil health and other environmental problems. A more robust and fundamental understanding of biological nitrogen fixation’s biochemistry and microbial physiology will enable strategies to promote new and more robust associations between nitrogen-fixing microorganisms and crop plants.

## Introduction

The availability of fixed nitrogen is of paramount importance to prototrophs, like plants. In agriculture, nitrogen fertilizers have become essential to maximizing crop yields to support the growing world population (1). Biological nitrogen fixation (BNF) is the reduction of atmospheric dinitrogen (N_2_) to ammonia (NH_3_) by diazotrophic bacteria and archaea, which accounts for ∼60% of the fixed nitrogen input into natural ecosystems (2). Nitrogenase, the enzyme catalyzing N_2_ reduction, is a significant energy sink as it requires large amounts of ATP and low potential electrons to produce NH_3_. There are three types of nitrogenase, termed Mo-, V-, and Fe-only nitrogenases, reflecting the metal cofactors’ composition in N_2_ reduction catalysis (3–5). Bacteria that contain V- and Fe-only nitrogenase are not dependent on Mo availability in the environment (6). Despite the similar features shared by the three nitrogenases, they differ in their reaction stoichiometry (7, 8). Whereby Mo-nitrogenase is the most efficient, requiring a minimum of 8 low potential electrons and 16 MgATP to convert N_2_ to 2 NH_3_ (**eq 1**) *in vitro*, and V- and Fe-only nitrogenases have lower catalytic activities and different reaction stoichiometries, requiring more electrons and ATP for catalysis and producing more H_2_ relative to NH_3_.

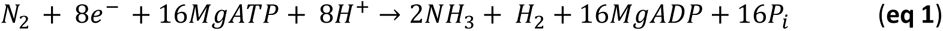

Diazotrophs, are physiologically diverse, including obligate aerobes, facultative anaerobes, anaerobic heterotrophs, anoxygenic or oxygenic phototrophs, and chemolithotrophs (9, 10). Under nitrogen-fixing conditions, diazotrophs must remodel their energy metabolism to provide nitrogenase with ATP and low potential electrons while protecting the enzyme from irreversible inactivation by oxygen (11). Oxygen protection is not an issue for strict anaerobes; however, the energy demands of nitrogen fixation during anaerobic metabolisms, such as fermentation, are profound relative to energy production per unit carbon (9). In contrast, oxygen respiration and photosynthesis can generate more energy for diazotrophic growth but protecting nitrogenase from oxygen inactivation becomes paramount. Diazotrophs that live in the air deal with protecting nitrogenase from inactivation through various mechanisms that involve conditionally, temporally, or spatially separating oxidative phosphorylation or photosynthesis from nitrogen fixation (12).

The ubiquitous soil bacterium *Azotobacter vinelandii* is arguably the most robust and productive nitrogen-fixing organism known (13, 14). *A. vinelandii* possesses a greater capacity to fix nitrogen than many other diazotrophs because of its ability to fix nitrogen under high oxygen concentrations. This ability is dependent on multiple mechanisms to protect nitrogenase from inactivation by oxygen (11, 15–17). One of the primary mechanisms involves harnessing a robust and dynamic respiratory metabolism to balance the high energy demands of nitrogen fixation while simultaneously consuming a high amount of oxygen at the membrane. This process, termed respiratory protection, maintains high enough respiration rates to sustain low oxygen tensions in the cytoplasm (11). A branch of the electron transport chain increases oxygen consumption by partially decoupling ATP synthesis from O_2_ consumption (14, 19). To supply energy for respiratory protection, *A. vinelandii* catabolizes sugars through the Entner-Doudoroff and pentose phosphate pathways to produce acetyl-CoA, then predominately uses the TCA cycle to deliver NADH (20, 21). During diazotrophic growth, *A. vinelandii* must efficiently balance the reduction of low potential electron carriers, ATP production using oxidative phosphorylation, and protection of the nitrogenase enzyme from oxygen through its dynamic electron transport system (ETS) (Fig 1).

**Figure 1.**
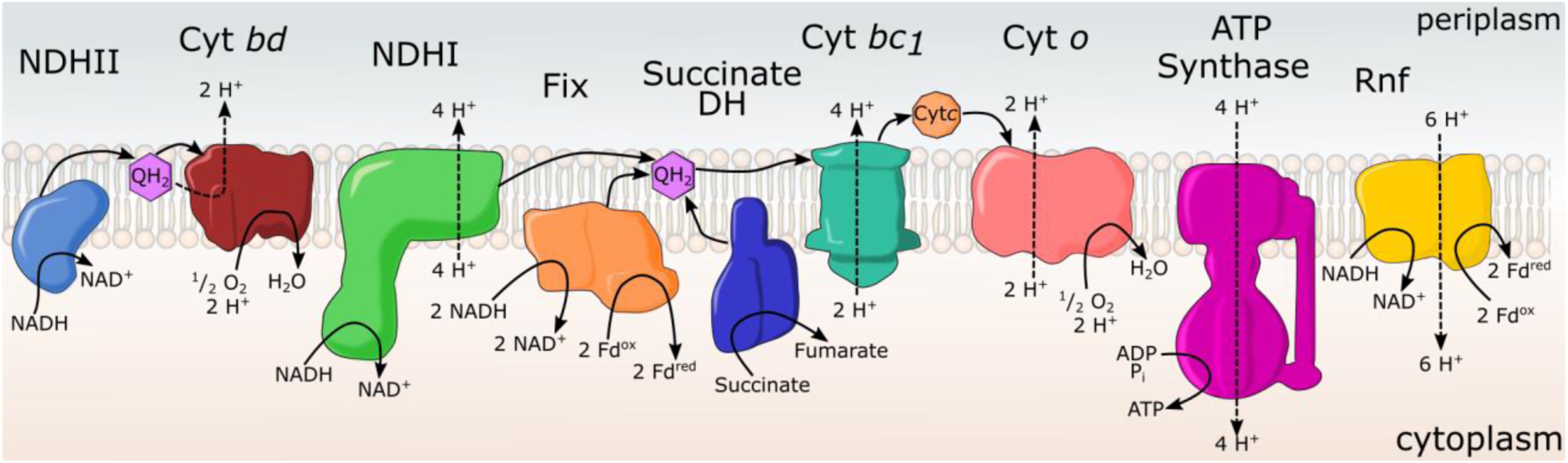
The ETS of *A. vinelandii* can be split into two paths. The first (left) is a respiratory protection pathway consisting of an uncoupled type II NADH dehydrogenase (NDHII) and terminal oxidase cytochrome *bd* (Cyt *bd*). The other path is a traditional proteobacterial electron transport chain with full proton coupled complexes: NADH dehydrogenase (NDHI), succinate dehydrogenase (Succinate DH), cytochromes *bc1*/*o* (Cyt *bc1*/*o*). The production of reduced ferredoxin (Fd) for nitrogenase is also part of the ETS with Fix and Rnf complexes. These enzymes work in parallel to balance energy production and oxygen protection.

*A. vinelandii* adjusts respiration through a branched respiratory chain that includes multiple dehydrogenases and terminal oxidases. The chain’s two branches are classified as 1) the proton-coupled branch and 2) the partially-coupled respiratory protection branch (15) (Fig. 2). These are mediated by two distinct NADH:quinone redox reaction complexes (NDH). The first, NDHI, is coupled to the transmembrane proton potential and is mechanistically similar to complex I of mitochondria (22). However, the second, NDHII, is induced at high aeration conditions and carries out NADH oxidation without translocating protons across the membrane, thus decoupling oxygen consumption from ATP generation (19). Other dehydrogenases (DH) can also donate to the quinone pool, including malate DH, succinate DH, and hydrogenases. The first two of these DHs do not increase in expression under nitrogen-fixing conditions. However, uptake hydrogenases are known to recycle electrons from the H_2_ produced by nitrogenase into the quinone pool (23, 24).

**Figure 2.**
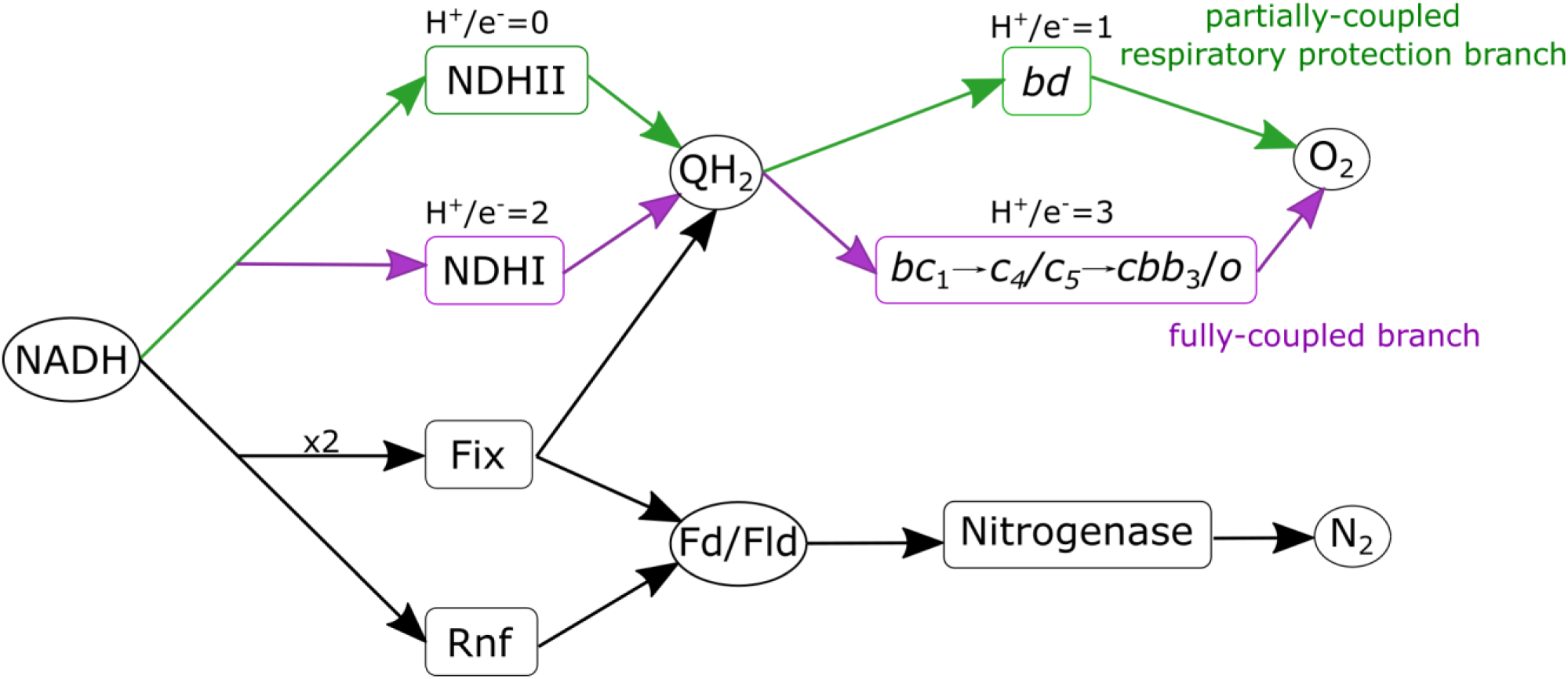
Paths of electrons in the electron transport system of *A. vinelandii.* NADH is potentially consumed by four different enzymes, uncoupled type II NADH dehydrogenase (NDHII), fully coupled NADH dehydrogenase (NDHI), Flavin based electron bifurcating Fix enzyme complex (Fix), and NADH:ferredoxin oxidoreductase (Rnf). There are two branches of the ETS that perform oxygen reduction: 1) partially-coupled respiratory protection branch (green) beginning with NDHII reducing quinone to quinol (QH_2_) and terminating with cytochrome *bd*, 2) fully-coupled branch (purple) beginning with NDHI reducing quinone, in turn reducing cytochrome *c* ending in cytochrome-*o* like terminal oxidase. Each branch translocate a different amount of protons per electron designated above the reaction names.

The oxidative side of the respiratory chain in *A. vinelandii* branches into several oxidases, including the *bc*1-complex, cytochrome *c*4/*c*5, and *o*-type or *cbb*_3_ terminal oxidases within the proton-coupled branch and cytochrome *bd*-type terminal oxidase within the partially-coupled respiratory protection branch (Fig. 2). Cytochrome *bd* accumulates under high aeration conditions, and knockout mutants lacking *bd* oxidase cannot grow diazotrophically at any aeration rate (13, 14, 25). The proton-coupled respiratory branch terminates in a classical cytochrome *bc1* reduction of cytochrome *c* to a terminal oxidase of cytochrome-*o* or *cbb*_3_ (26). This branch has not been as well-characterized in *A. vinelandii.* Still, kinetic evidence *in vivo* supports the existence of two cytochrome-c terminal oxidases (13).

Under nitrogen-fixing conditions, *A. vinelandii* directs most electrons to the reduction of NAD^+^, which has a reduction midpoint potential of ∼-320mV, while nitrogenase requires electrons with a lower potential of ∼-500mV (27). Additional energy is required to transfer electrons from NADH to lower potential electron carriers, such as ferredoxin (Fd) or flavodoxin (Fld). Under nitrogen-fixing conditions, *A. vinelandii* expresses membrane-associated Fix and Rnf complexes that catalyze the endergonic reduction of Fd/Fld by NADH (28, 29). Rnf uses the proton motive force to provide the additional energy required in the reaction (30–32). The Fix complex use flavin-based electron bifurcation in which Fix catalyzes the coordinated transfer of electrons from NADH to Coenzyme Q (CoQ) and Fd/Fld (29). The combination of branched electron transport to oxygen and the generation of Fd/Fld creates the ETS (Fig 2).

The metabolic energy cost of nitrogen fixation in *A. vinelandii* has recently been studied through carbon-based metabolomics (20) and investigated through multiple quantitative and metabolic models (33–35). However, these studies have not accounted for the dynamic *A. vinelandii’s* ETS and energy requirements, thus lack either insights into enzyme pathways of energy homeostasis or fail to predict growth under high oxygen and high substrate conditions. By integrating *A. vinelandii*’s energy metabolism dynamics under nitrogen-fixing conditions into the genome-scale metabolic model, an accurate growth and partitioning of resources can be predicted. Interestingly within the model, the carbon cost of aerobic nitrogen fixation is not entirely accounted for by the energy decoupling of the ETS’s partially-coupled respiratory protection branch. We show that under laboratory conditions of high carbon and high oxygen concentrations, large amounts of energy are dedicated to maintaining respiratory protection even in the presence of fixed nitrogen in the growth medium. Understanding the distribution of flux throughout the ETS is essential in the development of ammonia-excreting diazotrophs. The energy requirements and the metabolic bottlenecks for newly engineered ammonia-excreting strains may be predicted with the model.

The future of agriculture is dependent on an affordable, renewable, and environmentally sound supply of nitrogenous fertilizer. Synthetic biology and BNF have the potential of alleviating some dependency on traditional fertilizing techniques. Nevertheless, to maximize high throughput synthetic biology abilities, an accurate understanding of nitrogen fixation on the systems level is required. The metabolic model presented here is the first step in understanding some of the dynamics of this complex system.

## Results

### Curation of the metabolic model of A. vinelandii

Recently, a metabolic model (*iDT1278*) has been published that encompasses much of the *A. vinelandii* genome, establishing carbon and nitrogen sources using Biolog plate experiments (35). This model provides a framework for understanding the metabolism of *A. vinelandii* and is a valuable model for understanding the production of biopolymers. The model *iDT1278* lacked essential enzymes required for nitrogen fixation and failed to determine an accurate growth rate in standard laboratory conditions of complete aeration and at least 10g/L of sucrose or equivalent carbon (36). A new model (*iAA1300*) presented here builds off the model *iDT1278* by adding missing reactions and manually curating inadequately annotated constraints (Table S1). Key enzymes of the ETS were added to *iAA1300*, including Fix, NDHI, a quinone:cytochrome c oxidoreductase, V- and Fe-only nitrogenase, and a transhydrogenase, all of which have been biochemically or genetically determined to play a role during nitrogen fixation.

After manual curation, the model required central carbon metabolism constraints to represent experimental results more accurately. First, unlike other pseudomonads, *A. vinelandii* contains phosphofructose kinase (PFK) and has a complete Ember-Meyerhoff pathway (37). Nevertheless, multiple studies have shown that *A. vinelandii* utilizes the Entner-Doudoroff pathway (ED) (20, 38). Flux into the ED pathway and the glyoxylate shunt was constrained to a ratio determined previously by ^13^C-metabolic flux analysis (20). While these constraints directed carbon into the correct pathways, the predicted growth rate for model *iAA1300* was still inaccurate (Table 1).

**Table 1).** Growth rates and physiological parameters predicted form FBA results. All models have a glucose uptake rate of 15 mmol_glucose_/hr/gCDW. Entner-Doudoroff (ED), glyoxylate shunt (GS).

| Model | Growth rate |  |
| --- | --- | --- |
|  | Ammonia supplemented | Diazotrophic |
| Experimental | 0.27 | 0.22 |
| *iDT1278* | 1.87 | 1.28 |
| *iAA1300* | 1.43 | 0.98 |
| ED and GS Constrained | 1.37 | 0.93 |
| ED, GS, and Maintenance Constrained | 0.38 | 0.24 |

### Establishing parameters for accurate growth rate determination

Model *iAA1300* overestimated growth in almost every condition due to the lack of accurate non-growth associated maintenance flux (NGAM). Microbiologists since the ’50s have observed that the genus *Azotobacter* has an unusually high respiration rate leading to increased maintenance requirements (34–37). The high maintenance and respiration rate of *A. vinelandii* results in low biomass yields compared to other model proteobacteria. Quantitative modeling accurately described this phenomenon with high amounts of energy diverted to respiratory protection (33).

To translate the excess energy consumption into the genome-scale model, experimental data was used to predict an ATP maintenance (ATPM) rate under different O_2_ concentrations. Khula and Oelze (43) measured maintenance coefficients (mmol_Substrate_ · hr^−1^ · g of protein^−1^) of *A. vinelandii* growing in continuous diazotrophic cultures in different O_2_ concentrations and carbon sources using the Prit method (44) (Table 2). Maintenance coefficients increased as the O_2_ concentration increased in the bioreactor. Converting the experimentally determined maintenance coefficient to the genome-scale model ATPM (mmol_ATP_ · hr^−1^ · g CDW^−1^) requires an ATP/substrate ratio term. An issue arises with converting the maintenance coefficient to ATPM when considering the ATP produced per O_2_ consumed (P/O) ratio of the different branches of the ETS. The proton-coupled branch uses a mol of glucose to produce 32 mols of ATP, but the partially-coupled respiratory protection branch only produces 9 mols of ATP per mol of glucose. During high substrate and high O_2_ conditions, *A. vinelandii* requires decoupling of the ETS through the respiratory protection branch to maintain growth and minimize maintenance requirements (25, 45, 46).

**Table 2).**
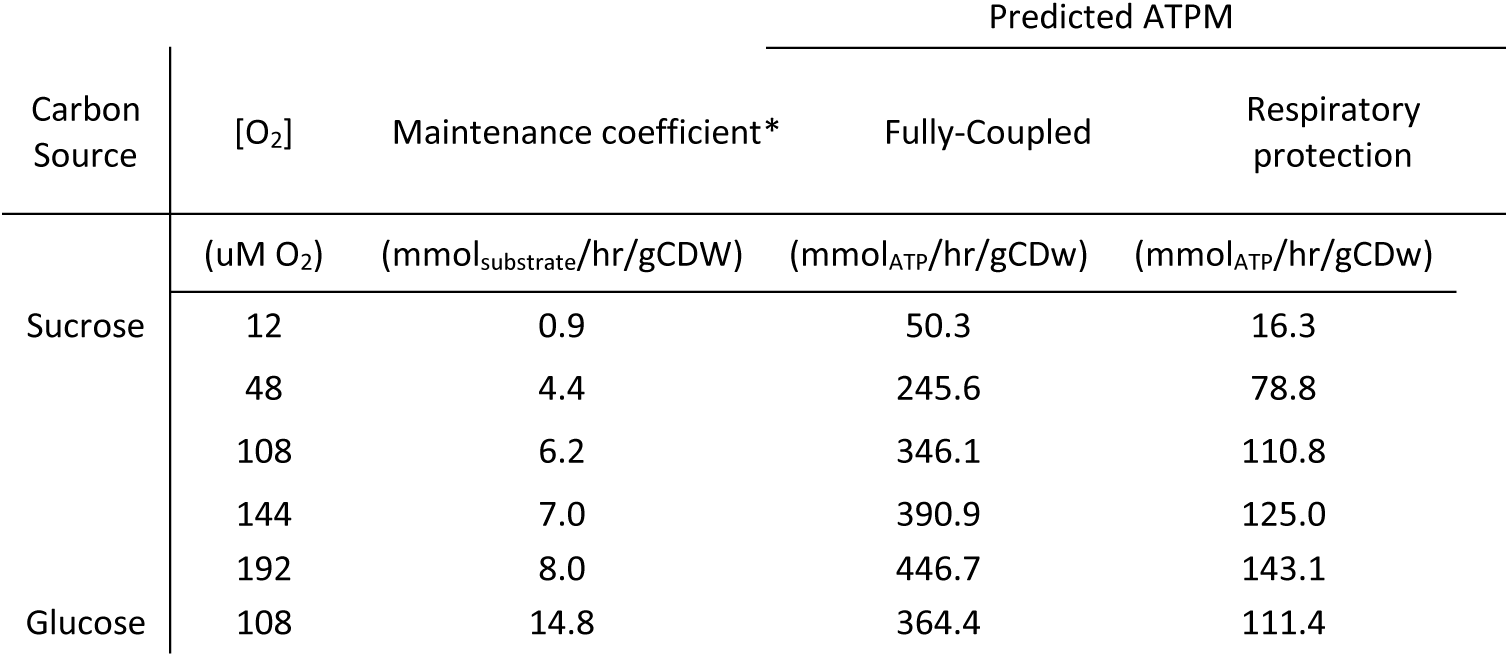
Experimental measured maintenance and predicted Non-growth associated maintenance (NGAM) for both branches of the ETC. *Maintenance coefficient from Kuhla and Oelze (43) converted from g of protein to gCDW.

To confirm the use of the partially-coupled respiratory protection branch, each path of the ETS network was tested to determine its accuracy to predict the growth rate. Two models were created, the first assuming all flux to O_2_ is directed through NDHI and cytochrome *co* (fully-coupled branch) and the second all flux to O_2_ through NDHII and cytochrome *bd* (respiratory protection branch) (Fig 2). In both models, substrate uptake rates were set to the experimentally determined maintenance coefficients for each O_2_ concentration (43). This uptake rate represents the substrate consumption when no growth occurs; therefore, all energy produced must go to NGAM. To determine the corresponding NGAM for each condition, flux through the reaction ATPM was increased until the growth rate reached zero (Table 2).

The model determined maintenance rates were then tested to predict growth rates at the different O_2_ concentrations. Each model was given the experimental substrate uptake rate and the predicted ATPM flux for each O_2_ concentration. Growth rates were then predicted and tested for error against experimental growth rates. Using the fully-coupled branch results in growth overestimates for all O_2_ concentrations (Fig. 3b). The respiratory protection branch model predicted growth rates with a minor error, especially for the lower O_2_ concentrations (Fig. 3a). The model is within reasonable error across all growth rates for O_2_ concentrations of 12, 48, and 108 µM (Table S2). For the higher O_2_ concentration of 144 and 192 µM, the respiratory protection branch model still overestimates growth. Showing a high O_2_ concentration respiratory protection and predicted maintenance could not compensate for total energy expenditure, requiring more ATPM flux than predicted.

**Figure 3).**
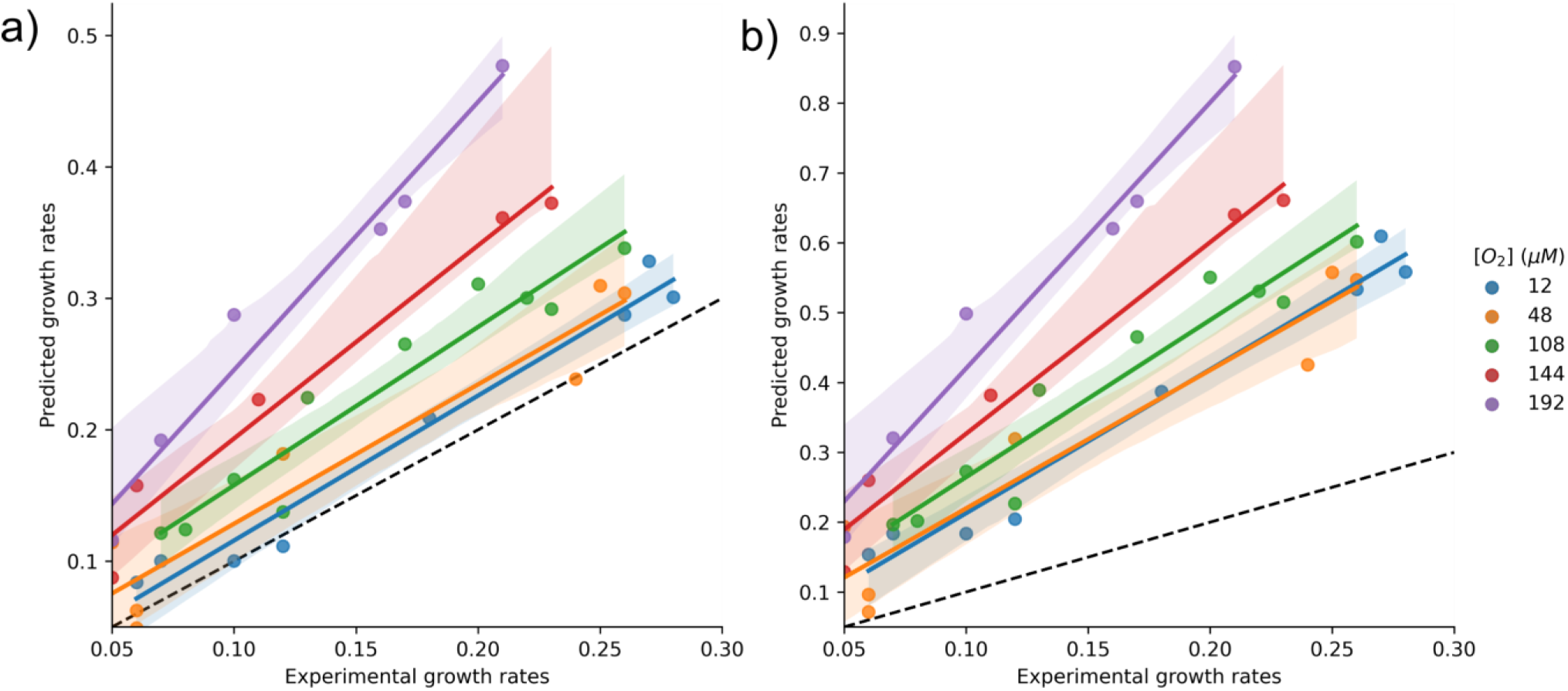
Comparison of theoretical growth rate and experimental growth rate for diazotrophic growth in different oxygen conditions. Tests the predictability of the for fully-coupled branch model and the respiratory protection branch model. Accurate prediction rates follow the x=y dotted line. Shaded color for 95% confidence interval for linear regression fit. A) Using the respiratory protection branch to determine ATPM flux gives accurate prediction of growth rate for lower oxygen concentrations of 12, 48, and 108 µM O_2_. Divergence from this trend occurs at 144 and 192 µM O_2_ where the model overestimates growth. B) Using the fully-coupled branch of the ETC to determine the ATPM flux cause an over estimation of growth in all conditions

### Assessment of growth yield in response to oxygen concentration

With detailed maintenance estimates, overall growth efficiencies can be further investigated. The growth yield was predicted using experimental sucrose uptake and plotted along with the experimentally determined growth yield (Fig. 4a). Similar to the growth rate predictions, the growth yield predictions indicate that the 12, 48, and 108 µM of O_2_ conditions are within error. In comparison, 144 and 192 µM of O_2_ are more challenging to predict with overestimating growth yields. The differentiation of growth yield between the 12 µM condition and higher O_2_ conditions initially seen in the experimental data can be reproduced with the model. The original paper of Kuhla and Oelze discussed this effect as the “decoupling of respiration” or respiratory protection (43). However, we have shown that the partially coupled respiratory protection branch is still required even at 12 µM of O_2_. To investigate this phenomenon more acutely, energy allocation during the increase of O_2_ concentration was plotted (Fig 3b). Both the flux to ATP synthase and O_2_ respiration (cytochrome *bd*) increase linearly with O_2_ concentration. However, partitioning of the ATP differentiates the 12 µM condition from the high O_2_ concentrations. The percentage of ATP consumed in ATPM reaction plateaus to around 60% of total ATP consumed for 48, 108, 144, and 192 µM of O_2_ while only at ∼ 30% for 12 µM of O_2_. This differentiation allows for more ATP to be utilized in biomass production, creating higher growth yields for the 12 µM of O_2_ concentration.

**Figure 4).**
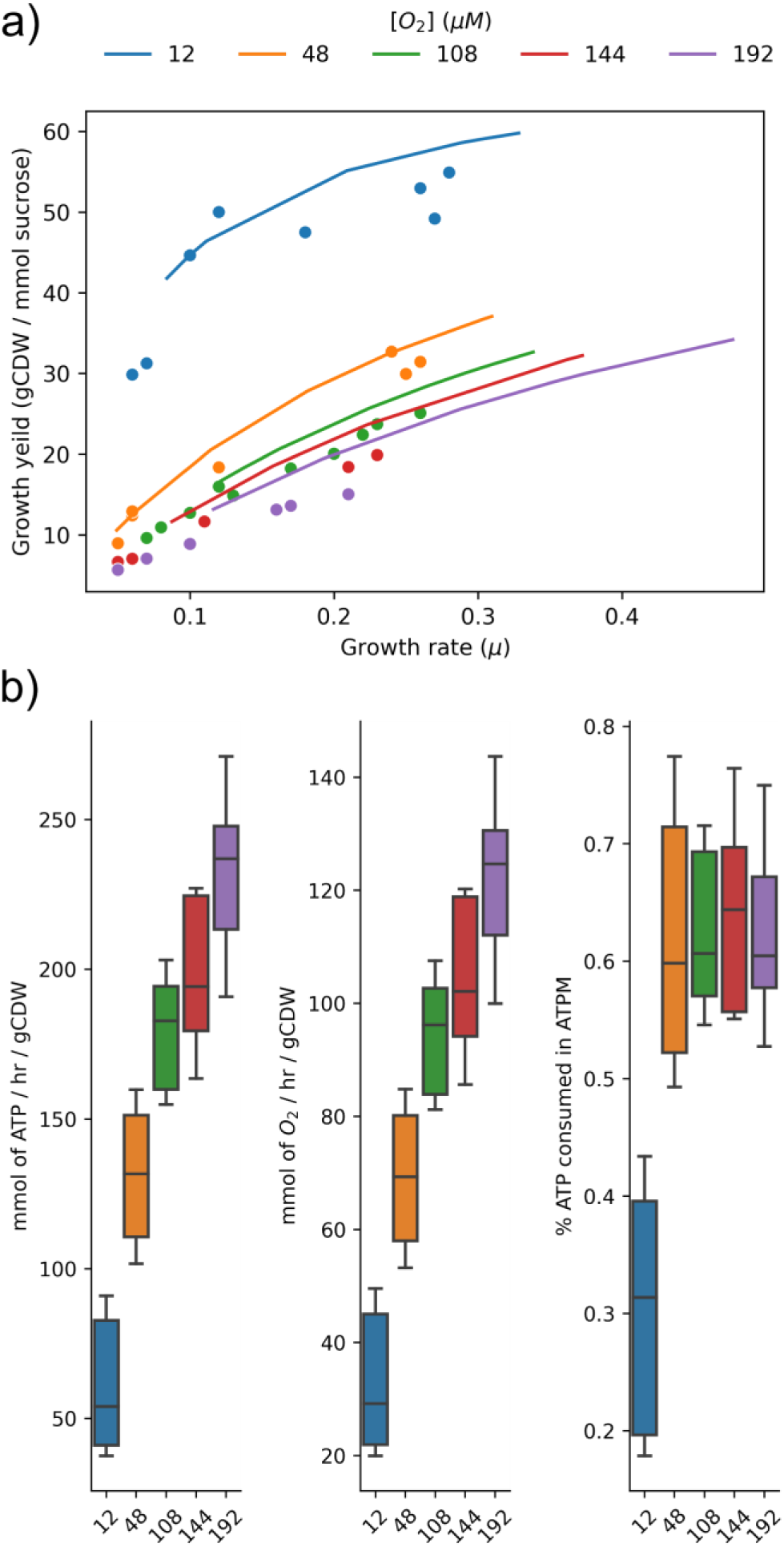
Allocation of carbon and energy under different O_2_ concentrations. A) Predicted growth yields were plotted for each O_2_ concentration across multiple growth rates (Lines). Plotted vs the experimental growth yields (points) show accurate predictions for lower oxygen concentrations. B) resource allocation across multiple O_2_ concentrations and growth rates, with flux through ATP synthase, respiration rates, and percentage of total ATP consumed in ATPM.

**Fig 5).**
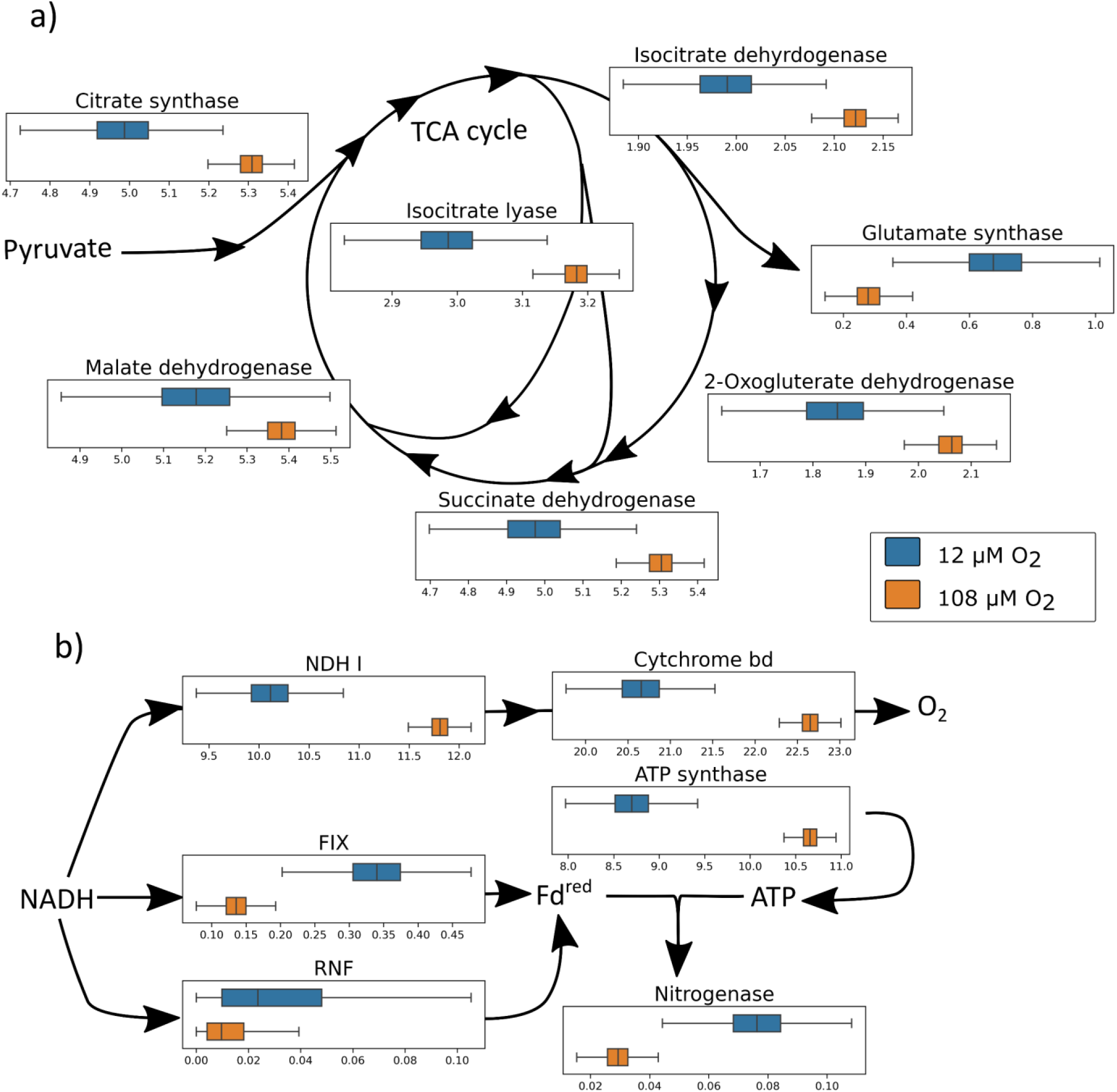
Histogram of flux samples normalized to sucrose uptake at different O_2_ concentrations. a) Key NADH producing enzymes of the TCA cycle show an increase of flux under higher [O_2_] (Orange) as compared to lower [O_2_]. While electron consuming reactions such as glutamate synthase reduces flux under high [O_2_]. b) The ETS under shifts fluxes to accommodate [O_2_] where the respiratory protection branch increases flux while electron flux to nitrogenase through ferredoxin (Fd^red^) is reduced. Flux is driven to oxygen reduction at the expense of nitrogen reduction. ATP generation is still maintained at a high rate to support NGAM in the model.

### Ammonia-assimilating conditions require high maintenance for accurate growth

The respiratory protection mechanism was considered a mechanism directly responsible for protecting nitrogenase from O_2_ damage (11). Experimentally, ammonia-supplemented conditions are similar to growth in diazotrophic conditions under high carbon, with comparable growth rates and biomass yield (20, 41). Modeling ammonia-supplemented growth shows the requirement of the partially coupled respiratory branch to minimize ATPM and accurately predict growth (Table 1). While there is a lack of accurate physiological details on *A. vinelandii* grown in high carbon and ammonia-supplemented media, similar respiration rates have been reported (41). The energy allocation under ammonia-supplemented growth shows that the ATP saved from nitrogen fixation is used for biomass production (Fig. S1).

### Effects of enzymes Rnf and Fix on accurate growth predictions

While the model shows the requirement of the respiratory protection branch to minimize flux through NGAM, little is known about Rnf and Fix’s roles during different O_2_ conditions. When determining ATPM flux, both branches of Rnf and Fix were considered, but either path did not affect the overall cost of NGAM. Under high O_2_ conditions, the percent of electron flux required for ferredoxin production is minimal compared to respiration. However, the different reaction mechanisms suggest these enzymes might play different roles within the ETS. This difference is accentuated when the uncoupled NADH dehydrogenase (NDHII) is used, and the energetic cost of Fix is not penalized. As the flux increases to nitrogenase and away from O_2_ reduction, Fix is favored as it can maintain a higher ATP production rate (Fig S2). Rnf can lower growth yields and increase O_2_ consumption, which could help predict high O_2_ concentrations more accurately.

### Flux sampling analysis reveals the dynamics of the ETS

A flux sampling approach was taken to further understand the network’s variability under high and low O_2_ conditions. While similar in concept to flux variability analysis, flux sampling analysis provides a range of all feasible solutions and allows for a distribution of the feasible fluxes, permitting statistics in determining shifts of change between conditions (47). To assess the effect of increased O_2_ during nitrogen fixation, the maintenance constraints defined in Table 2 under 108 and 12 µM of O_2_ were used. Flux balance analysis (FBA) showed similar growth rates of 0.202 hr^−1^ for 108 and 0.222 hr^−1^ for 12 µM of O_2_ making flux comparison approachable under these constraints. Samples were taken and normalized to sucrose uptake rate to compare electron allocation in the model. NADH production in the TCA increased in 108 µM of O_2_ compared to 12 µM of O_2_, but NADH consuming reactions such as glutamate synthase decreases flux in higher O_2_ concentrations, relative to carbon uptake (Fig 4a). The respiratory protection of the uncoupled NADH dehydrogenase and terminal oxidase cytochrome *bd* increases flux under higher O_2_ concentrations to protect nitrogenase and supply ATP for the increased maintenance rate. Electron transfer to nitrogenase through ferredoxin-reducing enzymes Rnf and Fix is reduced in higher O_2_ concentrations while ATP synthase is increased overall, leading to less flux to nitrogenase (Fig 4b).

### Efficient growth under metal limited conditions

*A. vinelandii* can adapt to the metal availability of its environment by using alternative nitrogenases. While the alternative nitrogenases use more common metals such as V and Fe, they are less efficient at reducing nitrogen than Mo-nitrogenase (**eq 2, 3**) (3). This inefficiency of ammonia production acts as an unnecessary sink for electrons, reducing the growth rate of *A. vinelandii* in Mo-limited conditions. Interestingly, while the cost to fix nitrogen rises 30% for V-nitrogenase and 60% for Fe-only nitrogenase, growth rates do not show an equal decrease (4, 48). Indicating a rearrangement of flux to compensate for the energy and electrons sink of the alternative nitrogenase while also maintaining O_2_ protection.

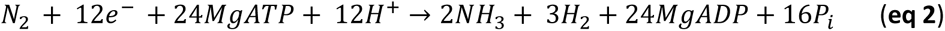

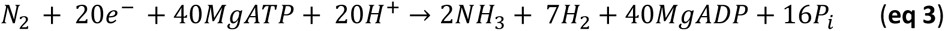

To demonstrate growth under alternative nitrogenase conditions, the flux through Mo-nitrogenase or both Mo- and V-nitrogenase was set to zero for V and Fe-only conditions, respectively. Growth rates were determined with FBA as well as O_2_ uptake rates and flux through each nitrogenase (Table 3). The model shows a slowing of growth by 13% for V-conditions and 31% for Fe-only conditions, which follows but is not proportional to the increased cost of nitrogenase turnover. Additionally, only a small increase of flux to O_2_ consumption is required to maintain energy production in alternative conditions.

**Table 3).** Growth rate and nitrogenase rates decrease as respiration increase as the model switches from Mo to V to Fe-only nitrogenase. All data from model with a sucrose uptake rate of 9 mmol_sucrose_/hr/gCDW and a ATP maintenance rate 110 mmol_ATP_/hr/gCDW. Units, grwoth rate (hr^−1^), O2 consumption (mmol_O2_/hr/gCDW), Nitrogenase flux (mmol_N2_/hr/gCDW).

|  | Mo | V | Fe |
| --- | --- | --- | --- |
| Growth rate | 0.22 | 0.17 | 0.13 |
| O<sub>2</sub> consumption | 98.16 | 99.40 | 101.14 |
| Nitorgenase flux | 1.04 | 0.91 | 0.72 |

To further investigate the rearrangement of the *A. vinelandii* metabolism to compensate for alternative nitrogenase flux, the flux sampling method was used to determine probabilities for flux changes between conditions. Flux samples were plot relative to the Mo-nitrogenase flux to better determine the alternative nitrogenases’ positive or negative effect (Fig S3). From the flux sampling, an increase in flux is seen through Fix as the alternative nitrogenases become less efficient, requiring more Fd. This effect is also true for uptake hydrogenase, which adapts to the increased hydrogen byproduct. The flux through the uncoupled NADH dehydrogenase is decreased as electron flux is compensated by hydrogenase and Fix (Fig S3).

### Optimal ammonia excretion under aerobic nitrogen-fixing conditions

Unlocking *A. vinelandii’s* nitrogen fixation regulatory system by deletion of *nifL* gene allows nitrogenase to be constitutively expressed even in the presence of high ammonia concentration in the media (36, 49). The ability to engineer an ammonia-excreting strain has been a target for genetic engineering for many decades. By simulating *A. vinelandii* to produce the maximum ammonia in agricultural or industrial scenarios, key insights can be developed for future engineering targets. To test the viability of ammonia excretion of the model and the effect of O_2_ maintenance, models of low and high O_2_ (12 and 108 µM of O_2_) were set to excrete ammonia at a rate of 3 mmol_Ammonia_ · hr^−1^ · g CDW^−1^ as estimated from Plunkett *et al*. (36). The increase of ammonia excretion essentially doubles the flux through nitrogenase and reduces the growth rate, respectively (Table 4). Ammonia-excreting strains start to excrete ammonia within the stationary phase during batch growths (36). In these conditions, cell growth would be minimal, and O_2_ would be limited due to cell density. As maximal ammonia excretion starts in the early stationary phase, the growth rate might not represent what is happening in the batch culture. Ammonia yields for high and low O_2_ are similar with 1.3 (mol_sucrose_/mol_ammonia_) predicted and while ∼1.4 (mol_sucrose_/mol_ammonia_) was experimentally determined. When O_2_ was increased to reduce the amount of time required for ammonia accumulation, an ammonia yield of ∼2.3 (mol_sucrose_/mol_ammonia_) was experimentally determined, and 3 (mol_sucrose_/mol_ammonia_) was predicted (36).

**Table 4).** Table of predicted fluxes for ammonia-assimilating, diazotrophic, and ammonia-excreting (ΔnifL) in high (108 µM) and low (12 µM) oxygen conditions. Growth rate is in gCDW of biomass / hr, all other fluxes are in mmol of metabolite/ gCDW / hr. Negative exchange rates are uptakes into the cell, while positive values are excretions.

| Conditions | Growth rate | Ammonia exchange | Respiration rate | Sucrose uptake | Nitrogenase Flux |
| --- | --- | --- | --- | --- | --- |
| Ammonia High O<sub>2</sub> | 0.30 | -3.19 | 95.24 | -9.00 | 0.00 |
| Ammonia Low O<sub>2</sub> | 0.33 | -3.48 | 34.07 | -4.00 | 0.00 |
| Diazotrophic High O<sub>2</sub> | 0.20 | 0.00 | 98.02 | -9.00 | 1.05 |
| Diazotrophic Low O<sub>2</sub> | 0.22 | 0.00 | 37.10 | -4.00 | 1.15 |
| *ΔnifL* High O<sub>2</sub> | 0.10 | 3.00 | 100.64 | -9.00 | 2.04 |
| *ΔnifL* Low O<sub>2</sub> | 0.12 | 3.00 | 39.71 | -4.00 | 2.14 |

## Discussion

The energy dynamics of aerobic metabolism and nitrogen fixation have been under discussion for many decades. The mechanism of respiratory protection first developed in *A. vinelandii* has been at the center of this discussion as this strategy has also been proposed for other aerobic nitrogen fixers such as oxygenic phototrophic cyanobacteria (50–52). Here we have presented a genome-scale metabolic model correctly estimating the effects of O_2_ on nitrogen fixation. The model has shown surprising results that contribute to more significant questions about aerobic metabolism during nitrogen fixation.

The decoupling of energy consumption and biomass accumulation combined with an exceptionally high respiration rate led to the proposal of respiratory protection (40, 53). This proposal was reinforced with the discovery of a branch of the ETS within *A. vinelandii* containing an uncoupled NADH dehydrogenase and cytochrome *bd* terminal oxidase with high Vmax and low affinity for O_2_ (11, 22, 25, 45, 54). Others have disagreed with the basic principles of respiratory protection as nitrogen fixation and O_2_ consumption are not correlated (15). Respiration rate plateaus after a concentration of 70 µM of O_2_ with only a corresponding slight decrease of nitrogenase rate (41). While these observations of plateauing of O_2_ respiration are valid, the decoupling of energy from biomass still increases with O_2_ concentration. Inomura et al. develop a quantitative mechanistic model showing increased respiratory protection, including maintenance as the O_2_ concentration increases (33). While the Inomura model accurately described respiratory protection and maintenance, an energy transfer efficiency parameter estimates the efficiency of the ETS to convert carbon into ATP. Using a stoichiometric model, we have provided evidence missing in other models and theories about respiratory protection.

Using experimental maintenance coefficients and the genome-scale model *iAA1300*, we have shown that the partially-coupled respiratory protection branch is required for all measured O_2_ concentrations. The partially coupled branch’s requirement is based on the assumption of minimizing NGAM within the model, which is high compared to other proteobacteria genome-scale models (55, 56). Minimization of NGAM is dependent on the decoupling of the ETS and the respiratory protection branch, but transcript expression and spectrographic data suggest that the fully-coupled branch may be active during normal nitrogen-fixing conditions (13, 23, 49, 57). More significant energy dissipation through the NGAM mechanism would be required if flux passes through both partially-coupled respiratory protection and fully-coupled branch.

O_2_ reduction and energy production decoupling are not entirely accounted for by the partially-coupled respiratory protection branch alone. The extra energy consumption required to maintain accurate growth is modeled as an ATP consumption reaction. This consumption is most likely many different reactions and does not have to be ATP, but two categories can be proposed 1) base metabolic reactions not accounted for in the model 2) reactions that respond to O_2_ and dissipate energy. For the first category, more accurate physiological data and biomass composition would help predict energetic needs. The current model predicts growth yields for 12, 44, 108 µM O_2_ concentrations, so a significant change in the biomass equation is not expected (Fig 4a). The *A. vinelandii* strain OP and derivatives such as strain DJ cannot produce alginate and do not produce Poly(3-hydroxybutyrate) under high O_2_ and continuous culture (58–61). However, energy-consuming mechanisms like protein turnover and unknown transport of metabolites or proteins might contribute to the basal NGAM. The second category of reactions responds to the O_2_ concentration and could be responsible for the energy dissipation. First, protein turnover and reactive O_2_ species in high O_2_ concentrations are unknown. Characterization of O_2_ sensitive *A. vinelandii* mutants showed only three of thirteen had decreased respiration or catalase rate, leaving mechanisms other than respiratory protection as possibly responsible for O_2_ sensitivity (62). Also, reactions known to be active during nitrogen fixation are challenging to model in steady-state such as proton leak, pili formation, and the *in vivo* stoichiometry of nitrogenase (23, 63, 64). Additionally, other reactions can consume O_2_ with a low enough reduction potential, including Mehler reactions or soluble terminal oxidases (65, 66).

Preserving high NGAM and the respiratory protection branch is also required for growth under ammonia supplemented conditions. While accurate data with high carbon and high ammonia is lacking, the diazotrophic maintenance rate predicted accurate growth rates for the ammonia supplement model. Under high sucrose and O_2_ concentrations, ammonia supplemented and nitrogen-fixing *A. vinelandii* respire at similar rates and offer similar steady-state protein levels (41), leading to the proposal that respiratory protection is not a mechanism for nitrogenase protection but a response to high carbon and O_2_ concentrations. The respiratory protection branch is regulated by cydR, an FNR regulatory protein that responds directly to O_2_ (25). The terminal oxidase cytochrome *bd* is not required for ammonia-supplemented growth (45). However, cytochrome-*d* deficient mutants grow poorly in ammonia-supplemented media if not inoculated at high cell density (45, 57). The decoupling of energy and high NGAM in ammonia supplemented growth could be maintained to keep the cytosol in low O_2_ or low redox potential for either reaction not related to nitrogenase or in preparation for nitrogenase expression.

To adapt to higher O_2_ concentrations, *A. vinelandii* must increase electron production. Flux sampling normalized to sucrose uptake shows an increased flux of energy-producing reaction of the TCA cycle and a decreased flux in other reactions such as glutamate synthase, Fix, Rnf, and nitrogenase. As the flux to O_2_ reduction and ATP generation increases, the percent of energy allocated to nitrogen-fixing reactions decreases. This explains why mutations in what should be necessary enzymes such as Fix or Rnf and uptake hydrogenase do not affect growth under standard high O_2_ conditions (24, 29, 67). Nevertheless, if more energy is allocated to nitrogenase under low O_2_ or Mo-limited conditions, these reactions become more critical. The increasing energy demand is significant during Mo-limited conditions, requiring 30% and 60% more energy for V-nitrogenase and Fe-only nitrogenase, respectively. Interestingly, *A. vinelandii* only grows slightly slower in media lacking Mo or lacking both Mo + V, under batch and continuous culture (48). The increased flux through hydrogenase and energy-conserving reactions like Fix allows *A. vinelandii* to maintain a higher growth rate. This general pattern shows when comparing Rnf’s energy-consuming proton motive force mechanism versus Fix’s energy-conserving electron bifurcation mechanism. As the cell moves away from the energy decoupling reaction, Fix can sustain growth. While kinetics and thermodynamics also influence the enzymes of the ETS, the stoichiometric pattern shows distinct roles for these enzymes.

Biological nitrogen fixation can alleviate the cost and damage caused by industrial nitrogenous fertilizer. Ammonia-excreting strain of *A. vinelandii* has supported plant growth and is a candidate for biofertilizer (68–70). Understanding the dynamics of metabolism under ammonia-excreting conditions will be essential to engineering more robust strains. Recent work optimized ammonia-excreting strains and showed up to 3 mmol of mmol_Ammonia_ · hr^−1^ · g CDW^−1^ excreted into the media (36). Modeling these rates shows a doubling of flux through nitrogenase and a halving of growth rate. Interestingly, ammonia-excreting strains grow at similar rates compared to WT, but the accumulation of ammonia in the media occurs in the stationary phase during batch culture. Suggesting that within WT nitrogenase flux limits growth in the log phase and is regulated in the stationary phase once carbon is low. In contrast, ammonia-excreting strains are also nitrogenase limited in the log phase but cannot regulate nitrogenase in the stationary phase. More dynamic modeling of this phenomenon will allow for more optimization and balance, leading to a technology that will maximize ammonia yield.

## Conclusion

We have been able to establish a genome-scale metabolic model of nitrogen fixation and adaptations to O_2_. This model gives a blueprint for future engineering strategies in nitrogen fixation and its ability to help offset nitrogenous fertilizer. We have shown that the nitrogen fixation model is affected by carbon concentration, O_2_ concentration, and ammonia supplementation. By adding the ETS to this model, we have discovered that the regulation of respiratory protection, which previously was proposed to be a mechanism for diazotrophic conditions, might be a general response to high carbon and high O_2_ conditions. The allocation of resources to an extraordinarily high maintenance rate is compensated by lowering growth yields and the rearrangement of the ETS. Future engineering in ammonia-excreting organisms must consider this balance between O_2_ reduction and nitrogen fixation and the complex relationship between the two.

## Materials and Methods

### Model curation

To build on top of the previous model, *iDT1278* (35), essential reactions for diazotrophic growth were corrected for stoichiometry and annotation or added to the model (Table S1). Enzymes of the ETS were added, including the electron bifurcating Fix complex, fully coupled NADH dehydrogenase I complex, cytochrome c oxidoreductase, nitrogenase homologs V-nitrogenase and Fe-only nitrogenase, as well as a soluble hydrogenase and a transhydrogenase. Other reactions were either reannotated or removed. All reactions using menaquinone were removed as *A. vinelandii* only contains quinone (71–73). Glucose uptake was constrained to reaction GLCt2pp (*gluP*) (74). The Rnf reaction stoichiometry was changed from 3 protons translocated to 6 protons translocated based on thermodynamic and kinetic analysis and found at the connected GitHub page in the RNF_stoich.ipynb jupyter notebook. The ED pathway and the glyoxylate shunt were constrained to ratios determined by metabolic flux analysis (20) using a custom python function based on COBRA MatLab function addRatioReaction (75).

The model was cleaned from dead-end reactions and orphaned metabolites while maintaining genome-relevant reactions. Out of the 2,289 reactions, 278 were essential, while 928 were categorized as blocked reactions where they could not carry flux. To allow the model to be built upon in the future, reactions with a corresponding gene are kept even if the reactions are blocked. Of the blocked reactions, 177 had no associated genes and were removed as they are not involved in gap-filling or gene homology. Following the removal of blocked “geneless” reactions, an additional 45 metabolites were also removed.

To determine the consistency and annotation standards within model *iAA1300,* memote software has been used with summary statistics reported in Table S3 and fully reports on the GitHub page (76).

### Flux balance analysis

All calculations were done with the cobrapy 0.21.0 (77). For flux balance analysis, the optimization problem is formulated as:

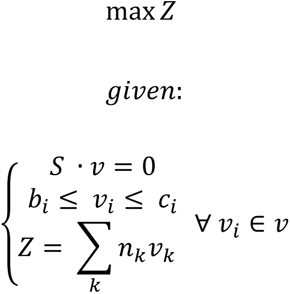

With Z being the biomass equation, with the stoichiometric coefficients *n_k_* and the biomass flux as *v_k_*. Biomass coefficients have a unit of hr^−1^ and represent the specific growth rate. *S* represents the stoichiometry matrix, and *v* is the flux vector. The scalars *b_i_* and *c_i_* are the lower and upper bounds for each flux *v_i_*.

### Maintenance rate quantification

Maintenance coefficients were taken from Table 1 in Kuhla and Oelze (43) and used as the sucrose uptake rate for the model. Under these conditions, the assumption is that all energy is going to NGAM, causing a zero growth rate. To determine the value of NGAM, the ATPM rate lower bound was increased until the growth reached zero, allowing all sucrose consumption to be allocated to NGAM. The determination of ATPM using this method depends on the ETS efficiency, so the fully-coupled branch and the respiratory protection branch of the ETS were used to determine a separated ATPM. Each ETS branch was then tested under different O_2_ concentrations giving a specific ATPM rate for experimentally determined maintenance coefficient (Table 2).

Testing the ATPM/NGAM values was done by setting the ATPM within the model and then increasing the sucrose uptake rate to the experimentally derived value found in Figure 4 of Kuhla and Oelze (43). Figure data points were taken using WebPlotDigitizer version 4.3 (78). With experimentally determined sucrose uptake rate and theoretically determined ATPM rates, a growth rate was predicted. The predicted growth rates were plotted against the known growth rates for both the fully-coupled and respiratory protection branches of the ETC. Determining mean standard error (MSE), mean absolute error (MAE), and root mean squared error (RSME) were all measured using the package scipy.stats (79).

### Flux sampling

Flux sampling analysis was conducted in COBRApy (77) using the optGpSampler (80) algorithm using 100000 samples with a thinning rate of 10000 in accordance with Hermann et al. (47). Model constraints for flux sampling were used from previous analysis for 108 µM and 12 µM O_2_ with the experimentally derived sucrose uptake rate of 9 and 4 mmol of sucrose hr^−1^ gCDW^−1^, respectively. The maintenance rates were used from the previous analysis of 110 mmol of ATP hr^−1^ gCDW^−1^ for 108 µM O_2_ and 16 mmol of ATP hr^−1^ gCDW^−1^ for 12 µM O_2_. Traditional FBA analysis was also performed to compare sampling analysis showing a growth rate of 0.202 hr^−1^ and 0.222 hr^−1^ for 108 µM and 12 µM O_2_, respectively. All plots were made in Python using Matplotlib.

### Ammonia excretion

The ammonia excreting model was determined using the glucose model based on constraints of Wu et al. (20) and ATPM rates determined above. The model was first tested for average growth under experimental conditions with an excretion rate of 3 mmol_Ammonia_ · hr^−1^ · g CDW^−1^ determined in Plunkett et al. (36).

### Alternative nitrogenases

The alternative nitrogenase enzymes of V-nitrogenase and Fe-only nitrogenase were tested for growth and flux sampling under standard sucrose conditions of 9 mmol of sucrose hr^−1^ gCDW^−1^ and an ATPM of 110 mmol of ATP hr^−1^ gCDW^−1^. While these conditions are not experimentally determined for alternative growth, they are a close approximation for batch growth cultures under high O_2_ and carbon but metal limited conditions. To determine growth rates, FBA was used, and flux sampling was conducted as stated above.

### Data Availability

Metabolic model *iAA1300* is attached in sbml format. All other data is available at https://github.com/alexander-alleman/Azotobactervinelandii_metabolicmodel. Metabolic models are saved in json and smbl format. All analysis and figure creation were documented in Jupyter notebooks.

---

**Figure S1).**
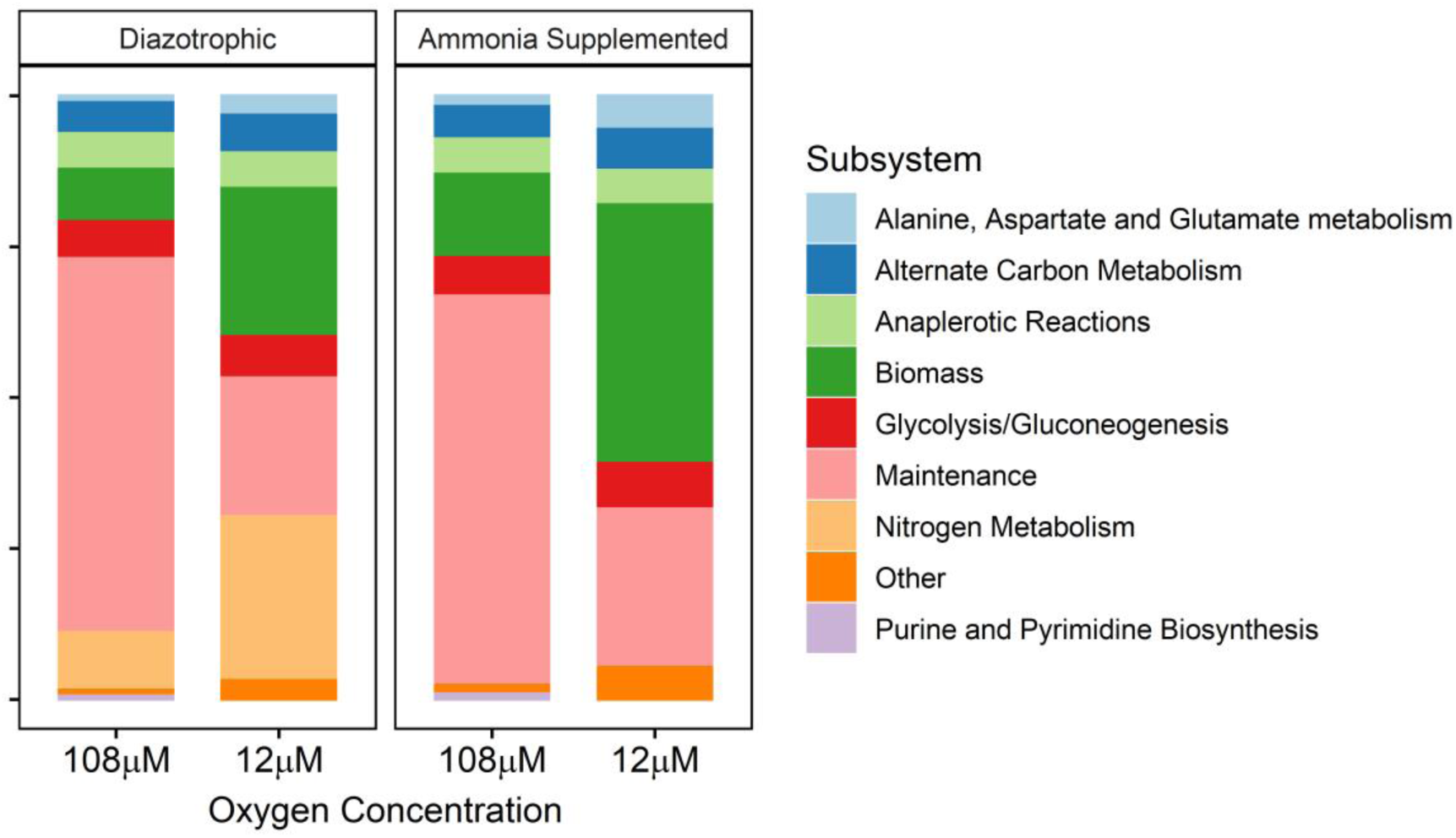
ATP allocation during high and low O_2_ concentration and with or without ammonia supplementation. If total flux of a subsystem was less than 1 mmol hr^−1^ gCDW^−1^ the subsystem was classified by “other”.

**Figure S2).**
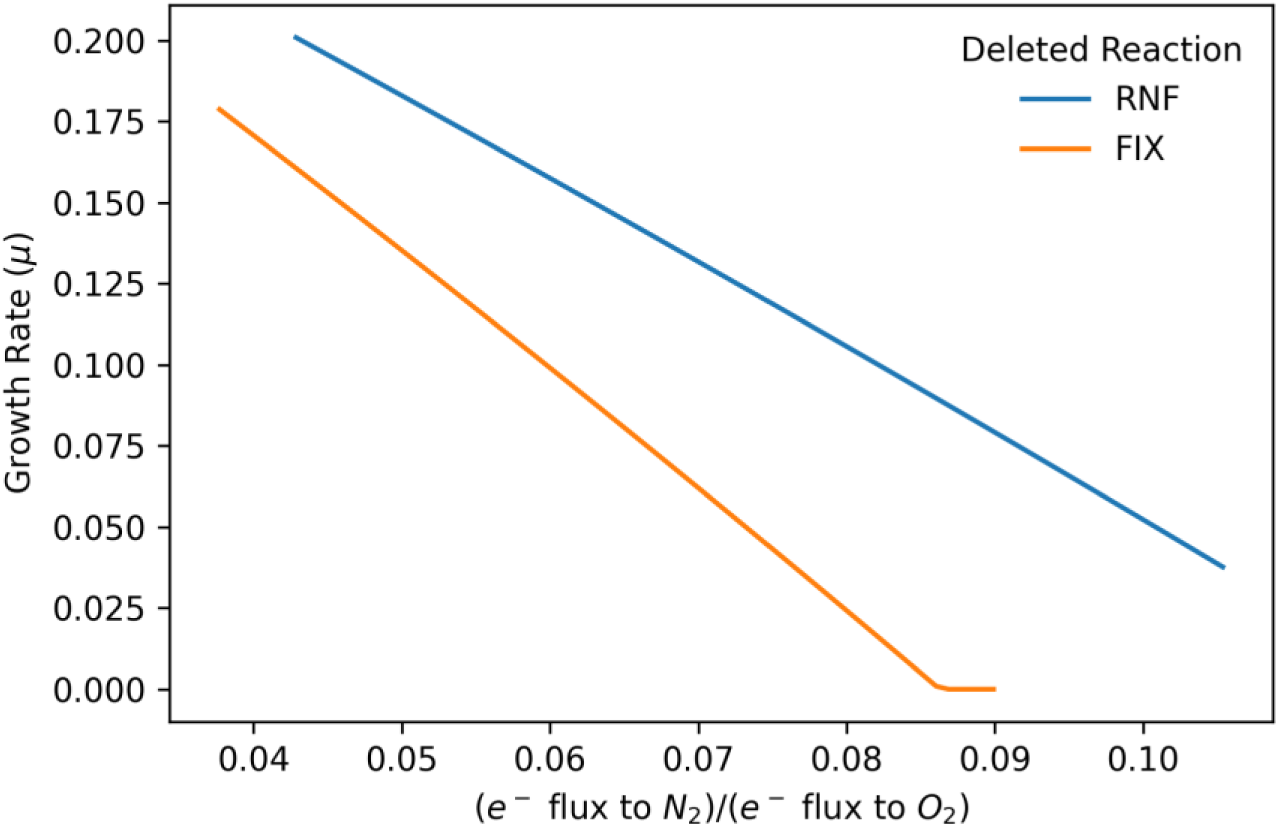
Growth rate vs flux to nitrogen reduction over flux to oxygen reduction. Models with either Rnf or Fix enzymes gene deletions were tested over a range of ratios of nitrogenase flux over terminal oxidase flux. As flux to nitrogenase is increased models without Fix is more susceptible and grows slower. Models without RNF can sustain a higher growth rate as flux to nitrogenase is increased.

**Figure S3).**
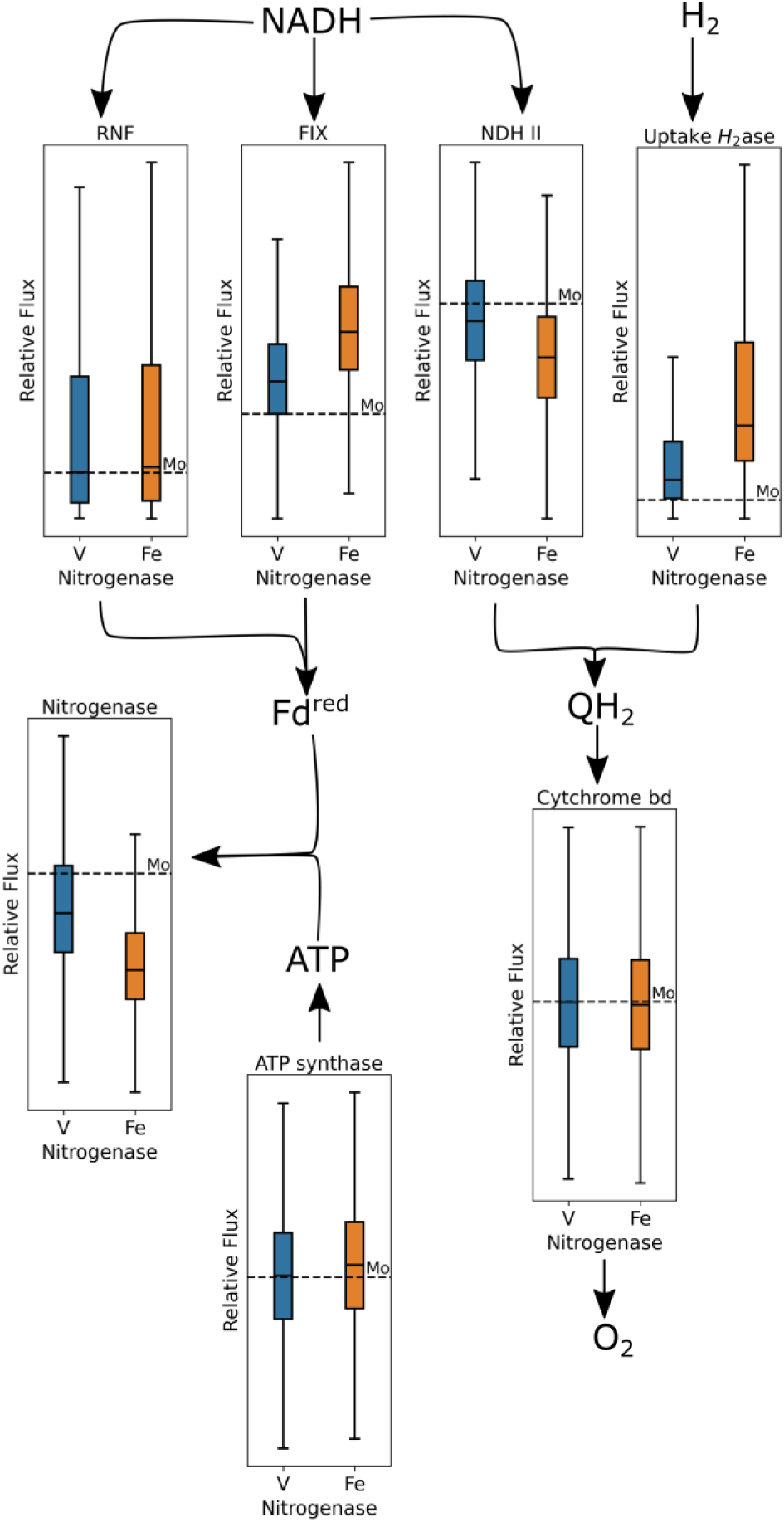
Flux sampling of the alternative nitrogenase models. Flux samples were normalized to oxygen uptake rates. To highlight shift form standard Mo-conditions each flux sample is then normalized to its corresponding Mo-flux. Dotted line is normalized Mo-condition flux or 1, so an increase in flux in other conditions are above this line or a decrease is below.

**Table S1).**
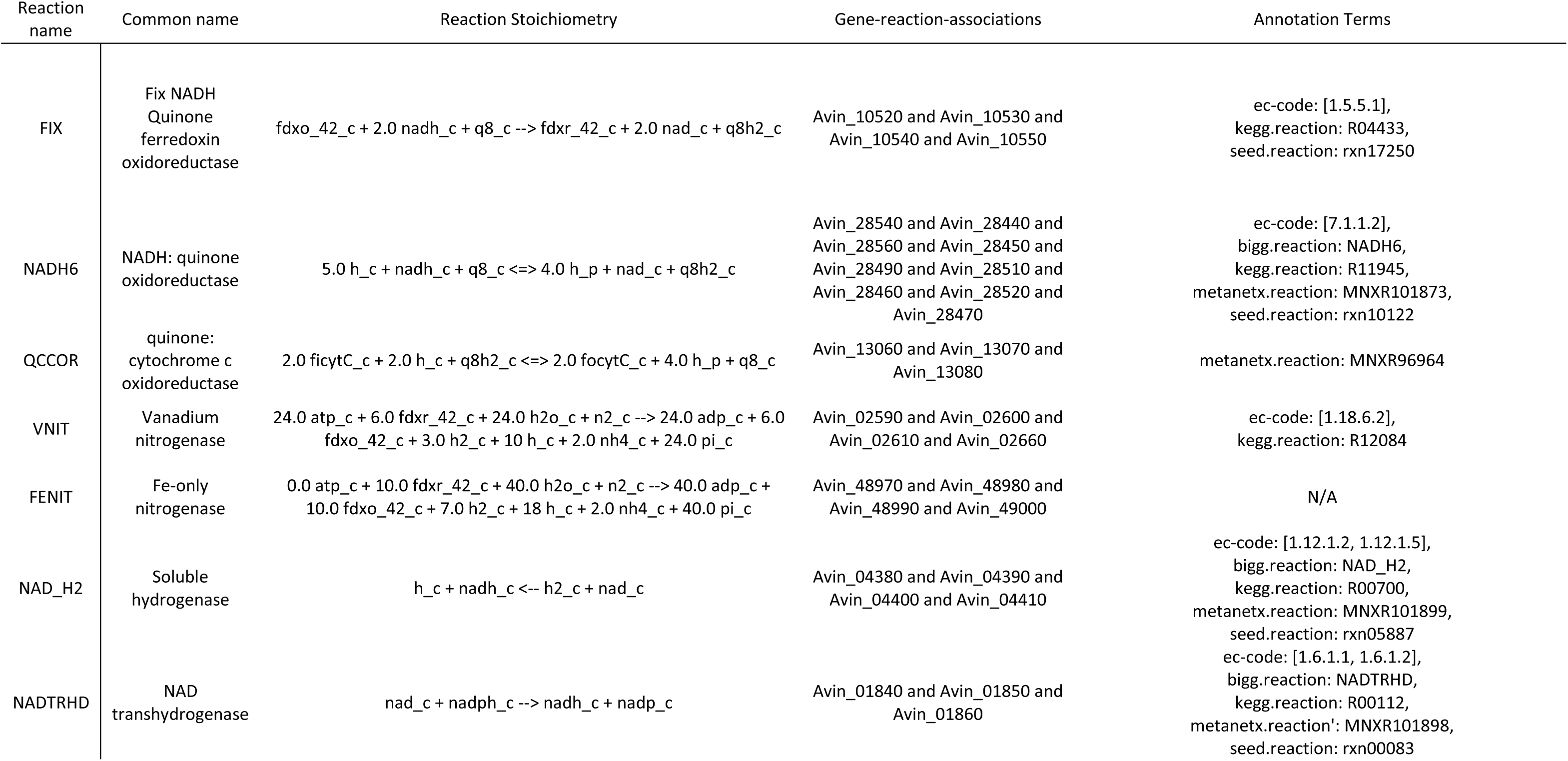
Reactions added to the model iAA1300 given common name, reaction stoichiometry and gene reaction associations. Annotation terms for FIX are terms for electron transfer flavoproteins (ETFs) as electron bifurcating enzyme complex is not yet in databases. V-nitrognease does have a kegg annotation but the stoichiometry is in accurate. Fe-only nitrogenase has no annotation in any database.

**Table S2).** The error of predicted growth rates compared to experimentally growth rates for both ETS branches under different oxygen concentrations. Mean square error (MSE), Mean absolute error (MAE), root mean squared error (RMSE).

| ETS Branch | $\mu\text{M O}_2$ | MSE | MAE | RMSE |
| --- | --- | --- | --- | --- |
| Respiratory Protection | 12 | 0.00 | 0.02 | 0.03 |
|  | 48 | 0.00 | 0.03 | 0.04 |
|  | 108 | 0.01 | 0.07 | 0.07 |
|  | 144 | 0.01 | 0.11 | 0.12 |
|  | 192 | 0.03 | 0.17 | 0.18 |
| Fully coupled | 12 | 0.05 | 0.19 | 0.22 |
|  | 48 | 0.04 | 0.17 | 0.21 |
|  | 108 | 0.07 | 0.25 | 0.26 |
|  | 144 | 0.11 | 0.29 | 0.32 |
|  | 192 | 0.20 | 0.41 | 0.44 |

**Table S3).** Basic model information and Memote criteria for the model presented in this paper (iAA1300) and the previous *A. vinelandii* model (iDT1278).

| Model | iAA1300 | iDT1278 |
| --- | --- | --- |
| Number of gene | 1300 | 1276 |
| Number of reactions | 2289 | 2469 |
| Number of metabolites | 1960 | 2003 |
| Metabolic coverage | 1.8 | 1.9 |
| Universally blocked reactions | 752 | 906 |
| Orphan Metabolites | 207 | 201 |
| Dead-end Metabolites | 191 | 190 |
| Memote criteria |  |  |
| Total score | 63% | 37% |
| Subtotal |  |  |
| Consistency | 86% | 85% |
| Annotation - Metabolites | 71% | 25% |
| Annotation - Reactions | 73% | 25% |
| Annotation - Genes | 0% | 0% |
| Annotation – SBO Terms | 46% | 0% |

## Acknowledgments

The authors would like to acknowledge Professor Bernd Markus Lange for valuable insight and editorial advice.

## References

1. Tittonell P, Giller KE. 2013. When yield gaps are poverty traps: The paradigm of ecological intensification in African smallholder agriculture. Field Crops Research 143:76–90.

2. Fowler D, Coyle M, Skiba U, Sutton MA, Cape JN, Reis S, Sheppard LJ, Jenkins A, Grizzetti B, Galloway JN, Vitousek P, Leach A, Bouwman AF, Butterbach-bahl K, Dentener F, Stevenson D, Amann M, Voss M. 2013. The global nitrogen cycle in the twenty-first century. Philosophical Transactions of the Royal Society B 368:1–13.

3. Mus F, Alleman AB, Pence N, Seefeldt LC, Peters JW. 2018. Exploring the alternatives of biological nitrogen fixation. Metallomics 10:523–538.

4. Bishop PE, Jarlenski DM, Hetherington DR. 1982. Expression of an Alternative Nitrogen Fixation System in *Azotobacter vinelandii*. Journal of bacteriology 150:1244–1251.

5. Chisnell JR, Premakumar R, Bishop PE. 1988. Purification of a second alternative nitrogenase from a *nifHDK* deletion strain of *Azotobacter vinelandii*. Journal of bacteriology 170:27–33.

6. Darnajoux R, Magain N, Renaudin M, Lutzoni F, Bellenger J-P, Zhang X. 2019. Molybdenum threshold for ecosystem scale alternative vanadium nitrogenase activity in boreal forests. PNAS 116:24682–24688.

7. Harris DF, Lukoyanov DA, Shaw S, Compton P, Tokmina-Lukaszewska M, Bothner B, Kelleher N, Dean DR, Hoffman BM, Seefeldt LC. 2017. Mechanism of N_2_ Reduction Catalyzed by Fe-Nitrogenase Involves Reductive Elimination of H_2_. Biochemistry 57:701–710.

8. Harris DF, Yang Z-Y, Dean DR, Seefeldt LC, Hoffman BM. 2018. Kinetic Understanding of N_2_ Reduction versus H_2_ Evolution at the E4(4H) Janus State in the Three Nitrogenases. Biochemistry 57:5706–5714.

9. Poudel S, Colman DR, Fixen KR, Ledbetter RN, Zheng Y, Pence N, Seefeldt LC, Peters JW, Harwood CS, Boyd ES. 2018. Electron transfer to nitrogenase in different genomic and metabolic backgrounds. Journal of Bacteriology 200:1–19.

10. Dos Santos PC, Fang Z, Mason SW, Setubal JC, Dixon R. 2012. Distribution of nitrogen fixation and nitrogenase-like sequences amongst microbial genomes. BMC Genomics 13:1–12.

11. Poole RK, Hill S. 1997. Respiratory protection of nitrogenase activity in *Azotobacter vinelandii*: roles of the terminal oxidases. FEMS microbiology reviews 17:303–317.

12. Bothe H, Schmitz O, Yates MG, Newton WE. 2010. Nitrogen Fixation and Hydrogen Metabolism in Cyanobacteria. Microbiol Mol Biol Rev 74:529–551.

13. D’Mello R, Hill S, Poole RK. 1994. Determination of the oxygen affinities of terminal oxidases in *Azotobacter vinelandii* using the deoxygenation of oxyleghaemoglobin and oxymyoglobin: Cytochrome *bd* is a low-affinity oxidase. Microbiology 140:1395–1402.

14. Bertsova YV, Bogachev AV, Skulachev VP. 1997. Generation of protonic potential by the *bd*-type quinol oxidase of *Azotobacter vinelandii*. FEBS Letters 414:369–372.

15. Oelze J. 2000. Respiratory protection of nitrogenase in Azotobacter species: Is a widely held hypothesis unequivocally supported by experimental evidence? FEMS Microbiology Reviews 24:321–333.

16. Sabra W, Zeng AP, Lünsdorf H, Deckwer WD. 2000. Effect of oxygen on formation and structure of Azotobacter vinelandii alginate and its role in protecting nitrogenase. Applied and Environmental Microbiology 66:4037–44.

17. Peña C, Peter CP, Büchs J, Galindo E. 2007. Evolution of the specific power consumption and oxygen transfer rate in alginate-producing cultures of *Azotobacter vinelandii* conducted in shake flasks. Biochemical Engineering Journal 73–80.

18. Lozano E, Galindo E, Peña CF. 2011. Oxygen transfer rate during the production of alginate by Azotobacter vinelandii under oxygen-limited and non oxygen-limited conditions 10:1–12.

19. Bertsova YV, Bogachev AV, Skulachev VP. 2001. Noncoupled NADH : ubiquinone oxidoreductase of *Azotobacter vinelandii* is required for diazotrophic growth at high oxygen concentrations. J Bacteriol 183:6869–6874.

20. Wu C, Herold RA, Knoshaug EP, Wang B, Xiong W, Laurens LML. 2019. Fluxomic Analysis Reveals Central Carbon Metabolism Adaptation for Diazotroph *Azotobacter vinelandii* Ammonium Excretion. 1. Scientific Reports 9:13209.

21. García A, Ferrer P, Albiol J, Castillo T, Segura D, Peña C. 2018. Metabolic flux analysis and the NAD(P)H/NAD(P)^+^ ratios in chemostat cultures of *Azotobacter vinelandii*. Microb Cell Fact 17.

22. Bertsova YV, Bogachev AV, Skulachev VP. 1998. Two NADH:ubiquinone oxidoreductases of *Azotobacter vinelandii* and their role in the respiratory protection. Biochimica et Biophysica Acta (BBA) - Bioenergetics 1363:125–133.

23. Hamilton TL, Ludwig M, Dixon R, Boyd ES, Dos Santos PC, Setubal JC, Bryant DA, Dean DR, Peters JW. 2011. Transcriptional profiling of nitrogen fixation in *Azotobacter vinelandii*. Journal of Bacteriology 193:4477–4486.

24. Noar J, Loveless T, Navarro-Herrero JL, Olson JW, Bruno-Bárcena JM. 2015. Aerobic hydrogen production via nitrogenase in *Azotobacter vinelandii* CA6. Applied and Environmental Microbiology 81:4507–4516.

25. Wu G, Cruz-Ramos H, Hill S, Green J, Sawers G, Poole RK. 2000. Regulation of cytochrome *bd* expression in the obligate aerobe *Azotobacter vinelandii* by *CydR* (Fnr). Sensitivity to oxygen, reactive oxygen species, and nitric oxide. Journal of Biological Chemistry 275:4679–4686.

26. Leung D, Oost J, Kelly M, Saraste M, Hill S, Poole RK. 1994. Mutagenesis of a gene encoding a cytochrome *o*-like terminal oxidase of *Azotobacter vinelandii* : A cytochrome *o* mutant is aero-tolerant during nitrogen fixation. FEMS Microbiology Letters 119:351–357.

27. Lanzilotta WN, Seefeldt LC. 1997. Changes in the midpoint potentials of the nitrogenase metal centers as a result of iron protein-molybdenum-iron protein complex formation. Biochemistry 36:12976–12983.

28. Boyd ES, Garcia Costas AM, Hamilton TL, Mus F, Peters JW. 2015. Evolution of molybdenum nitrogenase during the transition from anaerobic to aerobic metabolism. Journal of Bacteriology 197:1690–1699.

29. Ledbetter RN, Garcia Costas AM, Lubner CE, Mulder DW, Tokmina-Lukaszewska M, Artz JH, Patterson A, Magnuson TS, Jay ZJ, Duan HD, Miller J, Plunkett MH, Hoben JP, Barney BM, Carlson RP, Miller AF, Bothner B, King PW, Peters JW, Seefeldt LC. 2017. The electron bifurcating FixABCX protein complex from *Azotobacter vinelandii*: generation of low-potential reducing equivalents for nitrogenase catalysis. Biochemistry 56:4177–4190.

30. Hess V, Schuchmann K, Müller V. 2013. The ferredoxin: NAD^+^ Oxidoreductase (Rnf) from the acetogen *Acetobacterium woodii* requires Na^+^ and is reversibly coupled to the membrane potential. Journal of Biological Chemistry 288:31496–31502.

31. Curatti L, Brown CS, Ludden PW, Rubio LM, Kustu S. 2005. Genes required for rapid expression of nitrogenase activity in *Azotobacter vinelandii*. Proceedings of the National Academy of Sciences 102:6291–6296.

32. Biegel E, Schmidt S, González JM, Müller V. 2011. Biochemistry, evolution and physiological function of the Rnf complex, a novel ion-motive electron transport complex in prokaryotes. Cellular and Molecular Life Sciences 68:613–634.

33. Inomura K, Bragg J, Follows MJ. 2016. A quantitative analysis of the direct and indirect costs of nitrogen fixation: a model based on *Azotobacter vinelandii*. The ISME Journal 11:166–175.

34. Inomura K, Bragg J, Riemann L, Follows MJ. 2018. A quantitative model of nitrogen fixation in the presence of ammonium. PLOS ONE 13:e0208282.

35. Campos DT, Zuñiga C, Passi A, Del Toro J, Tibocha-Bonilla JD, Zepeda A, Betenbaugh MJ, Zengler K. 2020. Modeling of nitrogen fixation and polymer production in the heterotrophic diazotroph *Azotobacter vinelandii* DJ. Metabolic Engineering Communications 11:e00132.

36. Plunkett MH, Knutson CM, Barney BM. 2020. Key factors affecting ammonium production by an *Azotobacter vinelandii* strain deregulated for biological nitrogen fixation. Microbial Cell Factories 19:107.

37. Chavarría M, Nikel PI, Pérez-Pantoja D, de Lorenzo V. 2013. The Entner-Doudoroff pathway empowers *Pseudomonas putida* KT2440 with a high tolerance to oxidative stress: Perturbing the upper metabolism of *P. putida* with PFK. Environ Microbiol 15:1772–1785.

38. Wong TY, Yao X-T. 1994. The DeLey-Doudoroff Pathway of Galactose Metabolism in *Azotobacter vinelandii*. Applied and Environmental Microbiology 60:2065–2068.

39. Parker CA. 1954. Effect of Oxygen on the Fixation of Nitrogen by Azotobacter. 4408. Nature 173:780–781.

40. Dalton H, Postgate JR. 1968. Effect of Oxygen on Growth of *Azotobacter chroococcum* in Batch and Continuous Cultures. Microbiology, 54:463–473.

41. Post E, Kleiner D, Oelze J. 1983. Whole Cell respiration and nitrogenase activities in *Azotobacter vinelandii* growing in oxygen controlled continuous culture. Archives of Microbiology 134:68–72.

42. Parker CA, Scutt PB. 1960. The effect of oxygen on nitrogen fixation by Azotobacter. Biochimica et Biophysica Acta 38:230–238.

43. Kuhla J, Oelze J. 1988. Dependency of growth yield, maintenance and Ks-values on the dissolved oxygen concentration in continuous cultures of *Azotobacter vinelandii*. Archives of Microbiology 149:509–514.

44. Pirt SJ, Hinshelwood CN. 1965. The maintenance energy of bacteria in growing cultures. Proceedings of the Royal Society of London Series B Biological Sciences 163:224–231.

45. Kelly MJ, Poole RK, Yates MG, Kennedy C. 1990. Cloning and mutagenesis of genes encoding the cytochrome *bd* terminal oxidase complex in *Azotobacter vinelandii*: mutants deficient in the cytochrome *d* complex are unable to fix nitrogen in air. Journal of Bacteriology 172:6010–6019.

46. Kolonay JF, Maier RJ. 1997. Formation of pH and potential gradients by the reconstituted *Azotobacter vinelandii* cytochrome *bd* respiratory protection oxidase. Journal of bacteriology 179:3813–3817.

47. Herrmann HA, Dyson BC, Vass L, Johnson GN, Schwartz J-M. 2019. Flux sampling is a powerful tool to study metabolism under changing environmental conditions. 1. npj Systems Biology and Applications 5:1–8.

48. Natzke J, Noar JD, Bruno-Bárcena JM. 2018. *Azotobacter vinelandii* Nitrogenase Activity, Hydrogen Production, and Response to Oxygen Exposure. Applied and Environmental Microbiology 84:1–10.

49. Barney BM, Plunkett MH, Natarajan V, Mus F, Knutson CM, Peters JW. 2017. Transcriptional analysis of an ammonium excreting strain of *Azotobacter vinelandii* deregulated for nitrogen fixation. Applied and Environmental Microbiology 1–38.

50. Peschek GA, Villgrater K, Wastyn M. 1991. ‘Respiratory protection’ of the nitrogenase in dinitrogen-fixing cyanobacteria. Plant Soil 137:17–24.

51. Fay P. 1992. Oxygen relations of nitrogen fixation in cyanobacteria. Microbiol Rev 56:340–373.

52. Stal LJ. 2017. The effect of oxygen concentration and temperature on nitrogenase activity in the heterocystous cyanobacterium Fischerella sp. 1. Scientific Reports 7:5402.

53. Phillips DH, Johnson MJ. 1961. Measurement of dissolved oxygen in fermentations. Journal of Biochemical and Microbiological Technology and Engineering 3:261–275.

54. Ackrell BAC, Jones CW. 1971. The Respiratory System of *Azotobacter vinelandii*. European Journal of Biochemistry 20:22–28.

55. Nogales J, Mueller J, Gudmundsson S, Canalejo FJ, Duque E, Monk J, Feist AM, Ramos JL, Niu W, Palsson BO. 2020. High-quality genome-scale metabolic modelling of *Pseudomonas putida* highlights its broad metabolic capabilities. Environmental Microbiology 22:255–269.

56. Feist AM, Zielinski DC, Orth JD, Schellenberger J, Herrgard MJ, Palsson BØ. 2010. Model-driven evaluation of the production potential for growth-coupled products of *Escherichia coli*. Metabolic Engineering 12:173–186.

57. D’Mello R, Purchase D, Poole RK, Hill S. 1997. Expression and content of terminal oxidases in *Azotobacter vinelandii* grown with excess NH_4_ are modulated by O_2_ supply. Microbiology 143:231–237.

58. Castillo T, Heinzle E, Peifer S, Schneider K, Pena C. 2013. Oxygen supply strongly influences metabolic fluxes, the production of poly(3-hydroxybutyrate) and alginate, and the degree of acetylation of alginate in *Azotobacter vinelandii*. Process Biochemistry 48:995–1003.

59. Díaz-Barrera A, Urtuvia V, Padilla-Córdova C, Peña C. 2019. Poly(3-hydroxybutyrate) accumulation by *Azotobacter vinelandii* under different oxygen transfer strategies. J Ind Microbiol Biotechnol 46:13–19.

60. Martínez-Salazar JM, Moreno S, Nájera R, Boucher JC, Espín G, Soberón-Chávez G, Deretic V. 1996. Characterization of the genes coding for the putative sigma factor AlgU and its regulators MucA, MucB, MucC, and MucD in *Azotobacter vinelandii* and evaluation of their roles in alginate biosynthesis. Journal of bacteriology 178:1800–1808.

61. Setubal JC, Dos Santos P, Goldman BS, Ertesvåg H, Espin G, Rubio LM, Valla S, Almeida NF, Balasubramanian D, Cromes L, Curatti L, Du Z, Godsy E, Goodner B, Hellner-Burris K, Hernandez JA, Houmiel K, Imperial J, Kennedy C, Larson TJ, Latreille P, Ligon LS, Lu J, Mærk M, Miller NM, Norton S, O’Carroll IP, Paulsen I, Raulfs EC, Roemer R, Rosser J, Segura D, Slater S, Stricklin SL, Studholme DJ, Sun J, Viana CJ, Wallin E, Wang B, Wheeler C, Zhu H, Dean DR, Dixon R, Wood D. 2009. Genome sequence of *Azotobacter vinelandii*, an obligate aerobe specialized to support diverse anaerobic metabolic processes. Journal of Bacteriology 191:4534–4545.

62. Iwahashi H, Hachiya Y, Someya J. 1991. Isolation and characterization of oxygen sensitive mutants of *Azotobacter vinelandii*. FEMS Microbiology Letters 77:73–78.

63. Haaker H, Klugkist J. 1987. The bioenergetics of electron transport to nitrogenase. FEMS Microbiology Letters 46:57–71.

64. Hoffman BM, Lukoyanov D, Yang ZY, Dean DR, Seefeldt LC. 2014. Mechanism of nitrogen fixation by nitrogenase : the next stage. Chemical Reviews 114:4041–4062.

65. Varghese F, Kabasakal BV, Cotton CAR, Schumacher J, Rutherford AW, Fantuzzi A, Murray JW. 2019. A low-potential terminal oxidase associated with the iron-only nitrogenase from the nitrogen-fixing bacterium *Azotobacter vinelandii*. Journal of Biological Chemistry 294:9367–9376.

66. Sarkar D, Landa M, Bandyopadhyay A, Pakrasi HB, Zehr JP, Maranas CD. 2021. Elucidation of trophic interactions in an unusual single-cell nitrogen-fixing symbiosis using metabolic modeling. PLOS Computational Biology 17:e1008983.

67. Linkerhägner K, Oelze J. 1995. Hydrogenase does not confer significant benefits to *Azotobacter vinelandii* growing diazotrophically under conditions of glucose limitation. Journal of Bacteriology 177:6018–6020.

68. Ambrosio R, Ortiz-Marquez JCF, Curatti L. 2017. Metabolic engineering of a diazotrophic bacterium improves ammonium release and biofertilization of plants and microalgae. Metabolic Engineering 40:59–68.

69. Danyal K, Inglet BS, Vincent KA, Barney BM, Hoffman BM, Armstrong FA, Dean DR, Seefeldt LC. 2010. Uncoupling nitrogenase: Catalytic reduction of hydrazine to ammonia by a MoFe protein in the absence of Fe protein-ATP. Journal of the American Chemical Society 132:13197–13199.

70. Mus F, Crook MB, Garcia K, Garcia Costas A, Geddes BA, Kouri E-DD, Paramasivan P, Ryu M-H, Oldroyd GED, Poole PS, Udvardi MK, Voigt CA, Ané J-M, Peters JW. 2016. Symbiotic nitrogen fixation and challenges to extending it to non-legumes. Applied and environmental microbiology 82:3698–3710.

71. Wong T-Y, Maier RJ. 1984. Hydrogen-Oxidizing Electron Transport Components in Nitrogen-Fixing *Azotobacter vinelandii*. J BACTERIOL 159:5.

72. Jurtshuk P, Bednarz AJ, Zey P, Denton CH. 1969. L-malate oxidation by the electron transport fraction of *Azotobacter vinelandii*. Journal of Bacteriology 98:1120–1127.

73. Jones CW, Redfearn E. Electron Transport in *Azotobacter vinelandii*. Biochimica et Biophysica Acta 113:467–481.

74. Quiroz-Rocha E, Moreno R, Hernández-Ortíz A, Fragoso-Jiménez JC, Muriel-Millán LF, Guzmán J, Espín G, Rojo F, Núñez C. 2017. Glucose uptake in Azotobacter vinelandii occurs through a GluP transporter that is under the control of the CbrA/CbrB and Hfq-Crc systems. 1. Scientific Reports 7:858.

75. Heirendt L, Arreckx S, Pfau T, Mendoza SN, Richelle A, Heinken A, Haraldsdóttir HS, Wachowiak J, Keating SM, Vlasov V, Magnusdóttir S, Ng CY, Preciat G, Žagare A, Chan SHJ, Aurich MK, Clancy CM, Modamio J, Sauls JT, Noronha A, Bordbar A, Cousins B, El Assal DC, Valcarcel LV, Apaolaza I, Ghaderi S, Ahookhosh M, Ben Guebila M, Kostromins A, Sompairac N, Le HM, Ma D, Sun Y, Wang L, Yurkovich JT, Oliveira MAP, Vuong PT, El Assal LP, Kuperstein I, Zinovyev A, Hinton HS, Bryant WA, Aragón Artacho FJ, Planes FJ, Stalidzans E, Maass A, Vempala S, Hucka M, Saunders MA, Maranas CD, Lewis NE, Sauter T, Palsson BØ, Thiele I, Fleming RMT. 2019. Creation and analysis of biochemical constraint-based models using the COBRA Toolbox v.3.0. 3. Nature Protocols 14:639–702.

76. Lieven C, Beber ME, Olivier BG, Bergmann FT, Ataman M, Babaei P, Bartell JA, Blank LM, Chauhan S, Correia K, Diener C, Dräger A, Ebert BE, Edirisinghe JN, Faria JP, Feist AM, Fengos G, Fleming RMT, García-Jiménez B, Hatzimanikatis V, van Helvoirt W, Henry CS, Hermjakob H, Herrgård MJ, Kaafarani A, Kim HU, King Z, Klamt S, Klipp E, Koehorst JJ, König M, Lakshmanan M, Lee D-Y, Lee SY, Lee S, Lewis NE, Liu F, Ma H, Machado D, Mahadevan R, Maia P, Mardinoglu A, Medlock GL, Monk JM, Nielsen J, Nielsen LK, Nogales J, Nookaew I, Palsson BO, Papin JA, Patil KR, Poolman M, Price ND, Resendis-Antonio O, Richelle A, Rocha I, Sánchez BJ, Schaap PJ, Malik Sheriff RS, Shoaie S, Sonnenschein N, Teusink B, Vilaça P, Vik JO, Wodke JAH, Xavier JC, Yuan Q, Zakhartsev M, Zhang C. 2020. MEMOTE for standardized genome-scale metabolic model testing. 3. Nat Biotechnol 38:272–276.

77. Ebrahim A, Lerman JA, Palsson BO, Hyduke DR. 2013. COBRApy: COnstraints-Based Reconstruction and Analysis for Python. BMC Systems Biology 7:74.

78. Rohatgi A. WebPlotDigitizer User Manual Version 4.3 23.

79. Virtanen P, Gommers R, Oliphant TE, Haberland M, Reddy T, Cournapeau D, Burovski E, Peterson P, Weckesser W, Bright J, van der Walt SJ, Brett M, Wilson J, Millman KJ, Mayorov N, Nelson ARJ, Jones E, Kern R, Larson E, Carey CJ, Polat İ, Feng Y, Moore EW, VanderPlas J, Laxalde D, Perktold J, Cimrman R, Henriksen I, Quintero EA, Harris CR, Archibald AM, Ribeiro AH, Pedregosa F, van Mulbregt P. 2020. SciPy 1.0: fundamental algorithms for scientific computing in Python. 3. Nature Methods 17:261–272.

80. Megchelenbrink W, Huynen M, Marchiori E. 2014. optGpSampler: An Improved Tool for Uniformly Sampling the Solution-Space of Genome-Scale Metabolic Networks. PLOS ONE 9:e86587.

